# Diverse GABAergic neurons organize into subtype-specific sublaminae in the ventral lateral geniculate nucleus

**DOI:** 10.1101/2020.05.03.073197

**Authors:** Ubadah Sabbagh, Gubbi Govindaiah, Rachana D. Somaiya, Ryan V. Ha, Jessica C. Wei, William Guido, Michael A. Fox

## Abstract

In the visual system, retinal axons convey visual information from the outside world to dozens of distinct retinorecipient brain regions and organize that information at several levels, including either at the level of retinal afferents, cytoarchitecture of intrinsic retinorecipient neurons, or a combination of the two. Two major retinorecipient nuclei which are densely innervated by retinal axons are the dorsal lateral geniculate nucleus (dLGN), which is important for classical image-forming vision, and ventral LGN (vLGN), which is associated with non-image-forming vision. The neurochemistry, cytoarchitecture, and retinothalamic connectivity in vLGN remain unresolved, raising fundamental questions of how it receives and processes visual information. To shed light on these important questions, we labeled neurons in vLGN with canonical and novel cell type-specific markers and studied their spatial distribution and morphoelectric properties. Not only did we find a high percentage of cells in vLGN to be GABAergic, we discovered transcriptomically distinct GABAergic cell types reside in the two major laminae of vLGN, the retinorecipient, external vLGN (vLGNe) and the non-retinorecipient, internal vLGN (vLGNi). Within vLGNe, we identified transcriptionally distinct subtypes of GABAergic cells that are distributed into four adjacent sublaminae. Using trans-synaptic viral tracing and *in vitro* electrophysiology, we found cells in each these vLGNe sublaminae receive monosynaptic inputs from the retina. These results not only identify novel subtypes of GABAergic cells in vLGN, they suggest the subtype-specific laminar distribution of retinorecipient cells in vLGNe may be important for receiving, processing, and transmitting light-derived signals in parallel channels of the subcortical visual system.

**The vLGN is organized into subtype-specific sublaminae which receive visual input:** The ventral lateral geniculate nucleus (vLGN) is part of the visual thalamus. It can broadly be separated into two structural domains or laminae, the external vLGNe (which receives retinal input) and the internal vLGNi (receives no retinal input). In this study, we describe subtypes of transcriptomically distinct GABAergic neurons that populate the vLGN and organize into discrete, adjacent sublaminae in the vLGNe. Taken together, our results show four subtype-specific sublaminae of retinorecipient neurons in vLGNe.

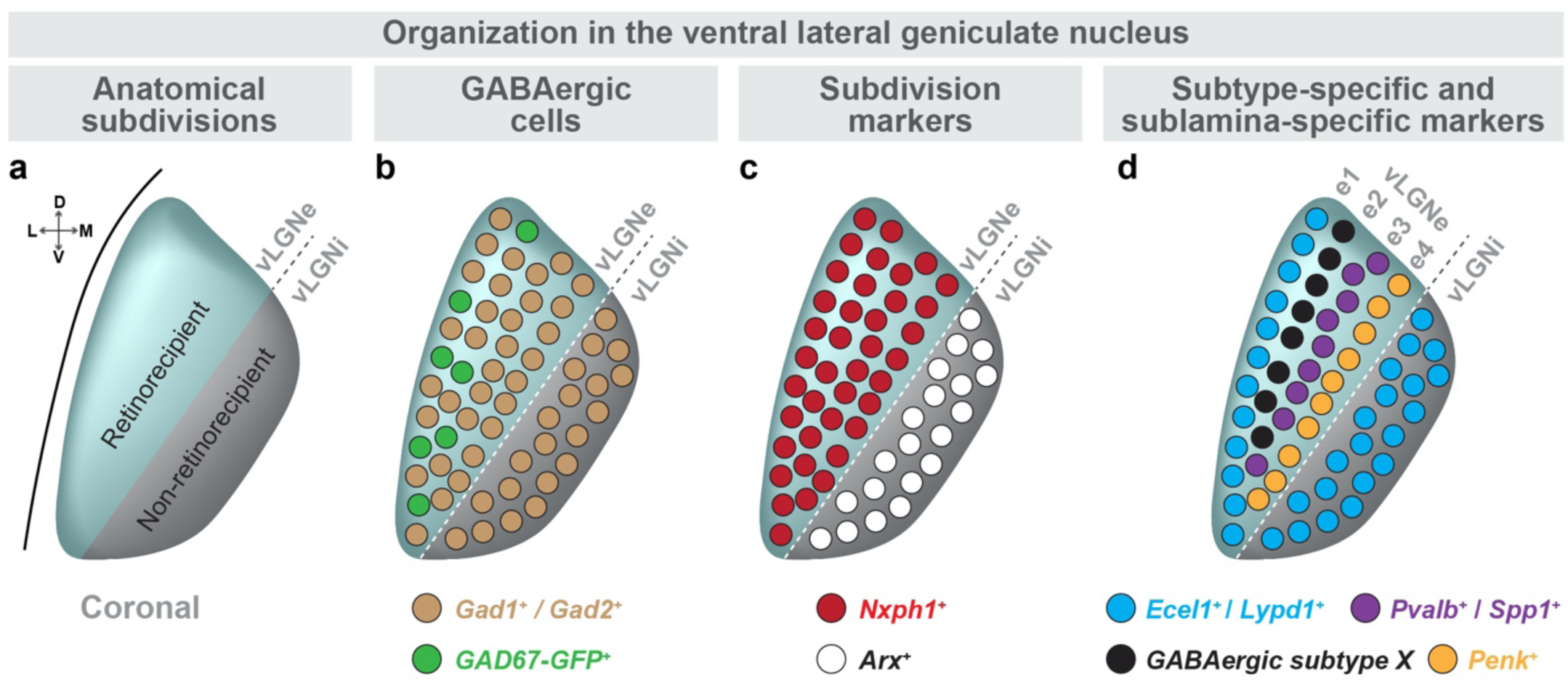

## Introduction

Information about the visual world is captured by the retina and transmitted by retinal ganglion cells (RGCs) to a diverse array of retinorecipient nuclei, including those in thalamic, hypothalamic, and midbrain regions (Fleming *et al.* 2006; Gaillard *et al.* 2013; Monavarfeshani *et al.* 2017; Morin & Studholme 2014; Martersteck *et al.* 2017). There is an organizational logic to these long-range retinal projections where RGCs, of which there are more than three dozen morphologically and functionally distinct subtypes, project to distinct and sometimes mutually exclusive retinorecipient regions (Hattar *et al.* 2006; Berson 2008; Dhande *et al.* 2011; Dhande *et al.* 2015; Kay *et al.* 2011; Osterhout *et al.* 2011; Yonehara *et al.* 2009). Many of these retinorecipient nuclei are critical to the execution of specific visual behaviors. For instance, retinal inputs to the dorsal lateral geniculate nucleus (dLGN) are important for image-formation and direction selectivity, those to the superior colliculus (SC) are important for gaze control, those to pretectal nuclei are important for pupillary reflexes and image stabilization, and those to the suprachiasmatic nucleus (SCN) are important for circadian photoentrainment (Dhande *et al.* 2015; Piscopo *et al.* 2013; Seabrook *et al.* 2017). Not only do RGCs project to different retinorecipient nuclei, but projections of distinct RGC subtypes are also segregated within a single retinorecipient region. For example, it has long been appreciated that projections from transcriptomically distinct ipsilateral and contralateral RGCs terminate in distinct domains of most rodent retinorecipient nuclei (Godement *et al.* 1984; Morin & Studholme 2014; Muscat *et al.* 2003; Jaubert-Miazza *et al.* 2005; Wang *et al.* 2016).

A long-standing objective of visual neuroscientists has been to characterize cell-type specific circuits in these retinorecipient regions, in terms of both inputs from RGCs and outputs to distinct downstream brain regions. For example, that distinct subtypes of RGCs terminate in different sublaminae of the SC (Dhande & Huberman 2014; Huberman *et al.* 2008; Huberman *et al.* 2009; Kim *et al.* 2010; Martersteck *et al.* 2017; Oliveira & Yonehara 2018). Post-synaptic to these retinal inputs are at least four morphologically and functionally distinct classes of retinorecipient neurons which are stellate, horizontal, wide-field, and narrow-field cells (Gale & Murphy 2014; Gale & Murphy 2016). Identifying subtype-specific retinocollicular circuitry facilitated the discovery that specific collicular cell types participate in different aspects of visually guided behavior (Hoy *et al.* 2019; Reinhard *et al.* 2019; Shang *et al.* 2018; Shang *et al.* 2015).

The dLGN, which processes and relays classical image-forming visual information to primary visual cortex, shares an organizational feature with SC in the kinds of retinal afferents it receives, where subtype-specific arborization of RGC axons been clearly characterized and forms so-called “hidden laminae” (Reese 1988; Martin 1986; Hong & Chen 2011). These hidden layers have been revealed by methods which individually label functionally and morphologically distinct classes of RGCs using transgenic reporter mouse lines (Huberman *et al.* 2008; Huberman *et al.* 2009; Kay *et al.* 2011; Kim *et al.* 2010; Kim *et al.* 2008). The dLGN is populated by just a few types of retinorecipient neurons, which include three classes of thalamocortical relay cells (X-like, Y-like, and W-like) and 1-2 classes of GABAergic interneurons (Arcelli *et al.* 1997; Jaubert-Miazza *et al.* 2005; Krahe *et al.* 2011; Leist *et al.* 2016; Ling *et al.* 2012). While their organization is not as ordered as their retinal afferents, classes of dLGN relay cells exhibit some regional preferences in their distribution, whereas interneurons are evenly dispersed throughout the nucleus (Krahe *et al.* 2011). Cell type-specific circuitry and function has also been demonstrated in dLGN, where W-like relay neurons receive input from direction-selective RGCs and in turn project to the superficial layers of mouse primary visual cortex (Cruz-Martín *et al.* 2014).

While our understanding of subtype-specific circuits has facilitated functional studies of SC and dLGN, there remain many retinorecipient regions about which such foundational information is unknown. One such region is the ventral LGN (vLGN), a portion of ventral thalamus that neighbors dLGN and is similarly innervated by retinal axons. Although less studied, it has been shown that vLGN is remarkably distinct from its dorsal counterpart in its transcriptome, proteome, cytoarchitecture, and circuitry (Harrington 1997; Su *et al.* 2011; Monavarfeshani *et al.* 2018; Sabbagh *et al.* 2018). In fact, distinct subtypes of RGCs project to vLGN and dLGN, and the majority of dLGN-projecting RGC classes fail to send collateral axons into vLGN, despite having to pass by it (or through it) on the way to dLGN (Huberman *et al.* 2008; Huberman *et al.* 2009; Kim *et al.* 2008). Retinal axons that target vLGN terminate in a lateral subdivision known as the external vLGN (vLGNe), which is cytoarchitectonically distinct from the internal vLGN (vLGNi) which receives little, if any, retinal input (Gabbott & Bacon 1994; Harrington 1997; Niimi *et al.* 1963; Sabbagh *et al.* 2018). The identity of retinorecipient cells in vLGNe remains largely unknown, although it is likely to include GABAergic cells (Huang *et al.* 2019), which represent the prevalent type of neuron in vLGN (Gabbott & Bacon 1994; Harrington 1997; Inamura *et al.* 2011).

Here, to address these gaps, we sought to determine the cell-types populating the vLGN, and their connectivity to retinal afferents. We assessed vLGN neurochemistry and cytoarchitecture by labeling cells with canonical and novel cell type markers. We found a richly diverse and tightly organized cellular landscape in vLGN, where transcriptomically distinct cell types are distributed in laminar subdomains, which appear to receive monosynaptic inputs from the retina. Our findings not only identify a novel organization of retinorecipient cells in vLGN, they suggest this order may be important for receiving, processing, and transmitting distinct light-derived signals in parallel channels of the subcortical visual system.

## Materials and Methods

### Animals

Wild type C57BL/6 mice were obtained from Jackson Laboratory. We obtained the following mice from Jackson Laboratory: *Pvalb-Cre* (JAX #: 008069, RRID:IMSR_JAX:008069), *Gad2-Cre* (JAX #: 010802, RRID:IMSR_JAX:010802), *Sst-Cre* (JAX #: 028864, RRID:IMSR_JAX:028864), *Sun1-stop-GFP* (JAX #: 021039, RRID:IMSR_JAX:021039), *Thy1-STOP-YFP* (JAX #: 005630, RRID: IMSR_JAX:005630), *ROSA-stop-tdT* (JAX #: 007909, RRID:IMSR_JAX:007909) and *GAD67-GFP* (JAX #: 007673, RRID:IMSR_JAX:007673). Animals were housed in a temperature – controlled environment, in a 12 hr dark/light cycle, and with access to food and water ad libitum. Both males and females were used in these experiments. Genomic DNA was isolated from tails genotyping as previously described (Su *et al.* 2010) using the following primers: *yfp*: fwd-AAGTTCATCTGCACCACCG, rev-TCCTTGAAGAAGATGGTGCG; *cre*: fwd-TGCATGATCTCCGGTATTGA, rev-CGTACTGACGGTGGGAGAAT; *sun1*: fwd-CTTCCCTCGTGATCTGCAAC, mut_rev-GTTATGTAACGCGGAACTCCA, wt-rev: CAGGACAACGCCCACACA; *tdt*: fwd-ACCTGGTGGAGTTCAAGACCATCT, rev-TTGATGACGGCCATGTTGTTGTCC. Animals were maintained and experiments conducted in compliance with National Institutes of Health (NIH) guidelines and approved protocols of the Virginia Polytechnic Institute and State University Institutional Animal Care and Use Committee (IACUC). Unless otherwise stated, n= number of animals and a minimum of three age-matched wildtype (and, where transgenic reporters were used, of same genotype) animals were compared in all experiments described.

### Immunohistochemistry (IHC)

Mice were anesthetized using 12.5 μg/mL tribromoethanol and transcardially perfused with PBS and 4% paraformaldehyde (PFA; pH 7.4). Extracted brains were kept in 4% PFA overnight at 4°C, and then incubated for at least 48 h in 30% sucrose in PBS. Fixed tissues were embedded in Tissue Freezing Medium (Electron Microscopy Sciences, Hatfield, PA, USA) and cryosectioned at 30 μm sections on a Leica CM1850 cryostat. Sections were air-dried onto Superfrost Plus slides (Fisher Scientific, Pittsburgh, PA, USA) and were dried for 15 min before being incubated in blocking buffer (2.5% bovine serum albumin, 5% Normal Goat Serum, 0.1% Triton-X in PBS) for 1 h at room temperature (or at 22°C). Primary antibodies were diluted in blocking buffer at the following dilutions and incubated on tissue sections at 4°C overnight: Calb1 (Swant, CB-38a, 1:1000) and Pvalb (Millipore-Sigma, MAB1572, 1:1000). Sections were then washed three times PBS and incubated in anti-mouse or anti-rabbit fluorescently conjugated secondary antibodies (Invitrogen Life Technologies, RRID:SCR_008410) diluted in blocking buffer (1: 1000) for 1 h at 22°C. Tissue sections were then washed at least 3 times with PBS, stained with DAPI (1: 5000 in water), and mounted using Vectashield (Vector Laboratories, Burlingame, CA, USA).

### Riboprobe production

Riboprobes were generated as previously described (Su *et al.* 2010; Monavarfeshani *et al.* 2018). *Gad1* (Gad1-F: TGTGCCCAAACTGGTCCT; Gad1-R: TGGCCGATGATTCTGGTT; NM_001312900.1; nucleotides 1099-2081), *Lypd1* (Lypd1-F: AAGGGAGTCTTTTTGTTCCCTC; Lypd1-R: TACAACGTGTCCTCTCAGCAGT; NM_145100.4; nucleotides 849-1522), *Arx* (Arx-F: CTGAGGCTCAAGGCTAAGGAG; Arx-R: GGTTTCCGAAGCCTCTACAGTT; NM_007492.4; nucleotides 1830-2729), *Ecel1* (Ecel1-F: CGCGCTCTTCTCGCTTAC; Ecel1-R: GGAGGAGCCACGAGGATT; NM_001277925.1; nucleotides 942-1895), *Nxph1* (Nxph1-F: ATAGGACAGGGCTGTCACCTTA; Nxph1-R: TTACTGAGAACAAGCTCCTCCC; NM_008751.5; nucleotides 1095-1695), *Spp1* (Spp1-F: AATCTCCTTGCGCCACAG; Spp1-R: TGGCCGTTTGCATTTCTT; NM_001204201.1; nucleotides 309-1263), *Sst* (NM_009215.1; nucleotides 7-550), and *Penk* (Penk-F: TTCCTGAGGCTTTGCACC; Penk-R: TCACTGCTGGAAAAGGGC; NM_001002927.3; nucleotides 312-1111) cDNAs were generated using Superscript II Reverse Transcriptase First Strand cDNA Synthesis kit (#18064014, Invitrogen) according to the manufacturer manual, amplified by PCR using primers designed for the above gene fragments, gel purified, and then cloned into a pGEM-T Easy Vector using pGEM-T Easy Vector kit, (#A1360, Promega) according to the kit manual. Anti-sense riboprobes against target genes were synthesized from 5 µg linearized plasmids using digoxigenin-(DIG) or fluorescein-labeled uridylyltransferase (UTP) (#11685619910, #11277073910, Roche, Mannheim, Germany) and the MAXIscript in vitro Transcription Kit (#AM1312, Ambion) according to the kit manual. 5 µg of riboprobe (20 µl) were hydrolyzed into ~0.5 kb fragments by adding 80 µl of water, 4 µl of NaHCO3 (1 M), 6 µl Na2CO3 (1 M) and incubating the mixture in 60°C. RNA fragments were finally precipitated in ethanol and resuspended in RNAase-free water.

### In situ hybridization (ISH)

ISH was performed on 30μm thin cryosections as described previously (Sabbagh *et al.* 2018). Sections were first allowed to air dry for 1hr at room temperature and washed with PBS for 5 min to remove freezing media. They were then fixed in 4% PFA for 10 min, washed with PBS for 15 min, incubated in proteinase K solution for 10 min, washed with PBS for 5 min, incubated in 4% PFA for 5 min, washed with PBS for 15 min, incubated in acetylation solution for 10 min, washed with PBS for 10 min, incubated in PBS-diluted 0.1% triton for 30 min, washed with PBS for 40 min, incubated in 0.3% H_2_O_2_ for 30 min, washed with PBS for 10 min, pre-hybridized with hybridization solution for 1 hr, before being hybridized with heat-denatured riboprobes at 62.5°C overnight. Sections were then washed for 5 times in 0.2X SSC buffer at 65°C. Slides were then washed with TBS, blocked, and incubated with horseradish peroxidase-conjugated anti-DIG (#11426346910, Roche) or anti-fluorescent antibodies (#11207733910, Roche) overnight at 4°C. Lastly, bound riboprobes were detected by a tyramide signal amplification system (#NEL753001KT, PerkinElmer).

### Anterograde axon and mono-synaptic tracing

Intravitreal injection of cholera toxin subunit B (to trace retinal terminals) was performed as previously described (Monavarfeshani *et al.* 2018; Su *et al.* 2011). Briefly, mice were anesthetized with isoflurane, and 1 μl of 1 mg/ml fluorescently conjugated Alexa-647-CTB (Invitrogen, C34778) was injected with a fine glass pipette using a picospritzer. After 3 days, animals were sacrificed and transcardially perfused with PBS followed by PFA.

A similar intravitreal injection of AAV2/1-hSyn-Cre-WPRE-hGH (2.5 × 10^13^ GC/mL, here referred to as AAV-Cre) was used to monosynaptically label retinorecipient neurons in the vLGN. 1.2 μl of AAV-Cre virus was injected either monocularly or binocularly at an approximate 45° angle relative to the optic axis. pENN.AAV.hSyn.Cre.WPRE.hGH was a gift from James M. Wilson (Addgene viral prep #105553-AAV1; RRID:Addgene_105553). Animals were sacrificed and perfused with PFA as described above 6-10 weeks after injection.

### Transcriptomic analyses

RNA sequencing experiments on developing WT vLGN and dLGN was described and published previously (Monavarfeshani *et al.* 2018).

### In vitro slice preparation and whole cell recording

*In vitro* recordings were conducted on genetically labeled vLGN neurons using methods described previously (Hammer *et al.* 2014). Mice were anesthetized with isoflurane, decapitated and brains were rapidly immersed in an ice-cold, oxygenated (95% O2/5% CO2) solution containing the following (in mM): 26 NaHCO3, 234 sucrose, 10 MgCl2,11 glucose, 2.5 KCl, 1.25 NaH2 PO4, 2 CaCl2. Coronal sections (270um) containing dLGN and vLGN were cut on a vibratome and placed in a chamber containing artificial cerebral spinal fluid (ACSF; in mM: 126 NaCl, 2.5 KCl, 1.25 NaH2PO4, 2.0 MgCl2, 26 NaHCO3, 2 CaCl2, 2 MgCl2 and 10 glucose, saturated with 95% O2/5% CO2, pH 7.3) at 32°C for 30 min and then at room temperature. Individual slices were transferred to a recording chamber maintained at 32°C and perfused continuously at a rate of 2.5 ml/min with oxygenated ACSF. Borosilicate pipettes were pulled using a two-step puller (Narishige) and filled with a solution containing the following (in mM): 117 K-gluconate, 13 KCl,1 MgCl2, 0.07 CaCl2, 0.01 EGTA, 10 HEPES, 2 Na-ATP, and 0.4 Na-GTP (pH 7.3, 290 osmol/L). For all recordings, biocytin (0.5%, Sigma) was included in the internal solution for intracellular filling and 3-D neuron reconstruction using confocal microscopy (Charalambakis *et al.* 2019; El-Danaf *et al.* 2015; Krahe *et al.* 2011). The final tip resistance of filled electrodes was 6 to 8 MΩ.

Whole cell patch recordings were made in current and voltage clamp using an amplifier (Multiclamp700B, Molecular Devices), filtered at 3-10kHz, digitized (Digidata 1440A) at 20kHz and stored on computer. Pipette capacitance, series resistance, input resistance, and whole-cell capacitance were monitored throughout the recording session devices.

To examine the intrinsic membrane properties of vLGN neurons, the voltages responses triggered by current step injection (−120 to +200 pA, 20 pA pulses, 600 ms) were recorded at resting membrane levels. Synaptic responses were recorded in voltage clamp (holding potential of −70mV) and evoked by electrical stimulation of the optic tract (OT) using bipolar tungsten electrodes (0.5 MΩ; A-M Systems) positioned just below the ventral border of vLGN. OT stimulation consisted of a 10Hz (10 pulses) train delivered at an intensity (25 to 200 µA) that evoked a maximal response (Hammer *et al.* 2014; Jaubert-Miazza *et al.* 2005).

### Quantification and imaging

To quantify the GABAergic neurons as a percentage of total cells in vLGN and dLGN, we labeled and counted *Gad1*^*+*^ (by ISH), *Gad2*^*+*^ (by *Gad2-Cre::Sun1-Stop-GFP* transgenic reporter), and *GAD67^+^-GFP* (by *GAD67-GFP* transgenic reporter) neurons and divided by all cells counted in that section by DAPI counterstaining. To quantify the density of a given GABAergic subtype in the two main subdomains of vLGN, we labeled with the respective subtype marker (after intravitreal CTB injection) and counted cells in the retinorecipient (CTB^+^) vLGNe and non-retinorecipient (CTB^−^) vLGNi and normalized to the area of the respective vLGN subdomain. Areas were measured by manual outlining of the border of vLGNe or vLGNi using ImageJ software (version 1.52n, NIH). Boundaries of vLGN or dLGN were determined by DAPI counterstaining. All quantification was performed on three biological replicates, counting at least three vLGN sections per mouse.

To quantify the spatial distribution of cell-type marker expression across the entire vLGN, we developed a custom line scan script (khatScan) that runs in ImageJ which overlays the vLGN with equally spaced lines. We opted for this approach over manually drawing lines to avoid user bias. A brief summary of how this script works: to determine the curvature of the vLGN in a particular image, khatScan prompts the user to draw a line along the optic tract adjacent to the vLGN, then automatically draws lines of a set length and number guided by that curve and plots the signal intensity across the x coordinates of each line. These intensities can then be averaged to determine where there is a specific enrichment for that marker in the vLGN. All imaging for quantification was performed on a confocal Zeiss LSM 700 microscope at 20x magnification and 0.5 digital zoom.

## Results

### Identification of distinct subtypes of GABAergic cells in vLGN

We first determined precisely what proportion of vLGN cells were GABAergic. In the brain, GABAergic neurons express *Gad1* and/or *Gad2*, genes which encode glutamate decarboxylases (GAD67 and GAD65 respectively), the biosynthetic enzymes necessary for the production of the neurotransmitter GABA. We therefore took two complementary approaches to label GABAergic interneurons: we performed *in situ* hybridization (ISH) with riboprobes generated against *Gad1* mRNA and we crossed *Gad2-Cre* mice (in which the Cre recombinase is expressed under the control of *Gad2* promoter) to *Rosa-Stop-Sun1/sfGFP* mice to transgenically label *Gad2*-expressing cells with a GFP-tagged nuclear protein (Mo *et al.* 2015). Both approaches revealed a dramatic enrichment of GABAergic cells in vLGN compared to dLGN, as expected from previous studies (Figure 1a-i) (Langel *et al.* 2018; Yuge *et al.* 2011). We found that >25% of cells in vLGN were *Gad1*^*+*^ (Figure 1c). A slightly higher percentage of vLGN cells (~40%) were GFP^+^ in *Gad2-Cre::Sun1-Stop-GFP* mice (Figure 1f). This increased percentage could be attributable to the limitations of mRNA detection by ISH and therefore represents a more accurate picture of the overall population of GABAergic cells in vLGN. This same approach labeled less than 10% of cells in dLGN, a number in line with previous reports (Arcelli *et al.* 1997; Su *et al.* 2020; Evangelio *et al.* 2018).

**Figure 1.**
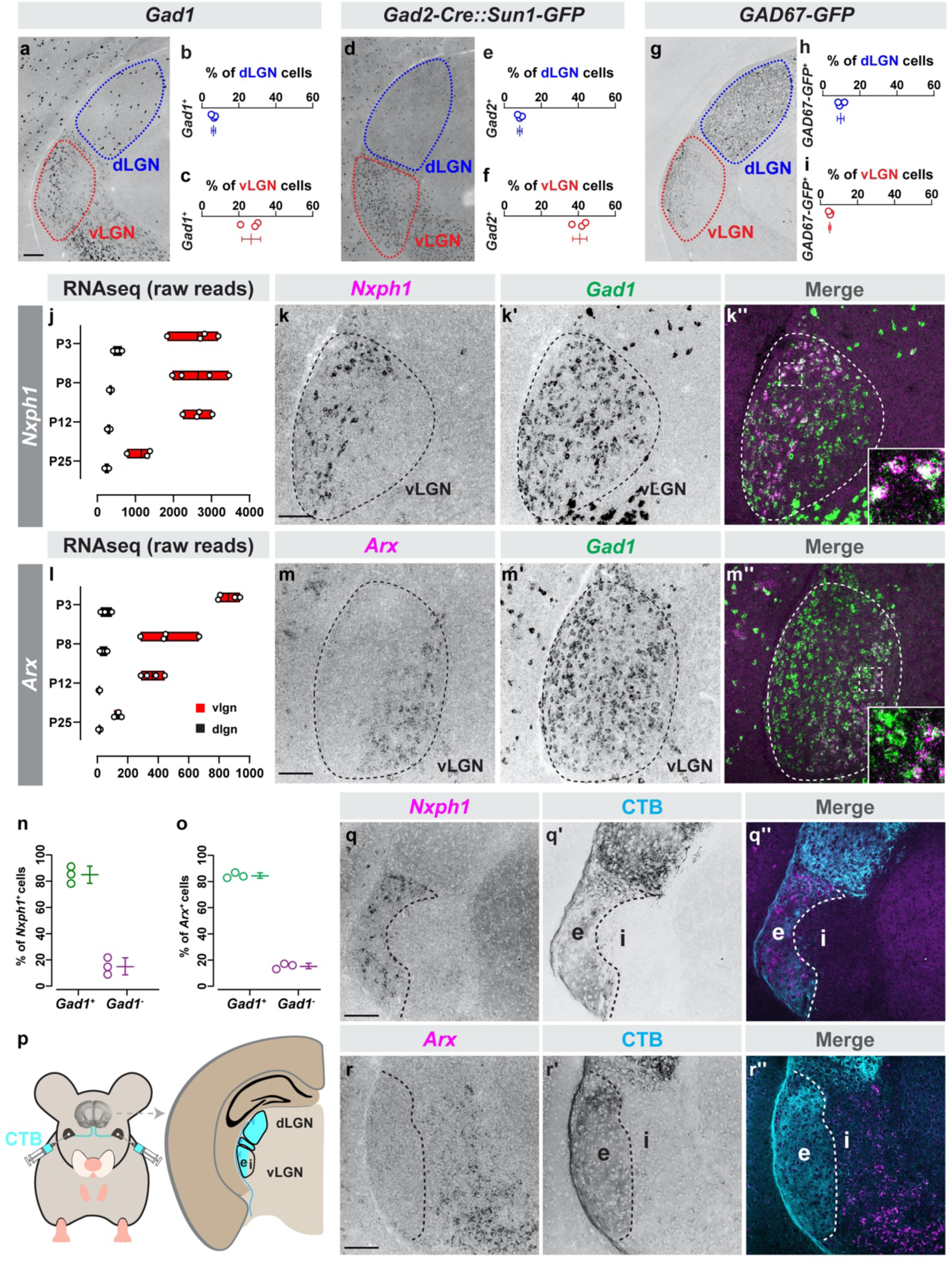
Cells in retinorecipient and non-retinorecipient zones of vLGN are molecularly distinct. (a) *In situ* hybridization for Gad1 mRNA in P60 dLGN and vLGN. (b,c) Quantification of the percentage of DAPI^+^ cells that are *Gad1*^*+*^ in dLGN (b) and vLGN (c). Data points represent biological replicates, bars represent mean ± SD. (d) Transgenic labeling of *Gad2+* GABAergic neurons in P60 LGN of *Gad2-Cre*::*Sun1-Stop-GFP* mice. (e,f) Quantification of the percentage of DAPI^+^ cells that are *Gad2+* in dLGN (e) and vLGN (f) of *Gad2-Cre*::*Sun1-Stop-GFP*. Data presented as in (c). (g) Transgenic labeling of GFP^+^ cells in P60 LGN of *GAD67-GFP* mice. (h,i) Quantification of the percentage of DAPI^+^ cells that are GFP^+^ in dLGN (h) and vLGN (i) of *GAD67-GFP* mice. Data presented as in (c). (j) Raw transcript reads of *Nxph1* mRNA in vLGN and dLGN obtained by RNAseq. Individual datapoints plotted as white circles, min/max values are confined to the red bars, and vertical black line with bars depicts mean. (k-k’’) Double ISH for *Nxph1* and *Gad1* mRNAs in P10 vLGN. (l) Raw transcript reads of *Arx* mRNA in vLGN and dLGN obtained by RNAseq, presented as in (j). (m-m’’) Double ISH in vLGN using riboprobes generated against *Arx* and *Gad1* mRNAs in P10 vLGN. (n,o) Quantification of the percent of *Nxph1+* (n) or *Arx*^*+*^ (o) cells that co-express *Gad1* mRNA. Data presented as in (c). (p) Intravitreal injection of Alexa-conjugated Cholera Toxin subunit B (CTB) labels retinothalamic projections. (q,r) ISH for *Nxph1* (q) and *Arx* (r) mRNAs in P10 CTB-labeled vLGN. All scale bars = 100 µm.

Several groups, including our own, have previously used a *GAD67-GFP* transgenic line to label most (if not all) GABAergic cells in dLGN (Charalambakis *et al.* 2019; Su *et al.* 2020; Seabrook *et al.* 2013). However few GABAergic neurons are labeled in vLGN of these mice (Figure 1g). In fact, we found less than 5% of cells in vLGN were GFP^+^ in *GAD67-GFP* (Figure 1i). The dramatically fewer GFP^+^ cells, compared to the number of GABAergic cells observed by labeling with either *Gad1* ISH or *Gad2-Cre::Sun1-Stop-GFP*, is due to the fact that the *GAD67-GFP* labels only a subset of thalamic GABAergic cells – likely local inhibitory interneurons – in visual thalamus (Su *et al.* 2020).

Together, these results suggested that multiple subtypes of GABAergic cells exist in vLGN, unlike in dLGN where GABAergic cells are relatively homogenous (Leist *et al.* 2016; Jager *et al.* 2016; Kalish *et al.* 2018). For this reason, we next asked whether we could identify novel molecular markers to characterize the heterogeneity of GABAergic neurons in vLGN. We assessed gene expression profiles in the developing mouse vLGN and dLGN in previously generated RNAseq datasets (Monavarfeshani *et al.* 2018; He *et al.* 2019). Our rationale was to identify candidate cell type makers by focusing our attention on genes which were: a) enriched in vLGN but not dLGN, b) expressed by GABAergic cells in other brain regions, and/or c) expressed with different developmental patterns which could indicate expression by different subsets of neurons. To characterize if any of these genes labeled distinct populations of neurons in vLGN, we generated >40 riboprobes to perform ISH (**Table S1**). We also took advantage of available cell-specific reporter mice and antibodies for this screen.

**Table S1.**
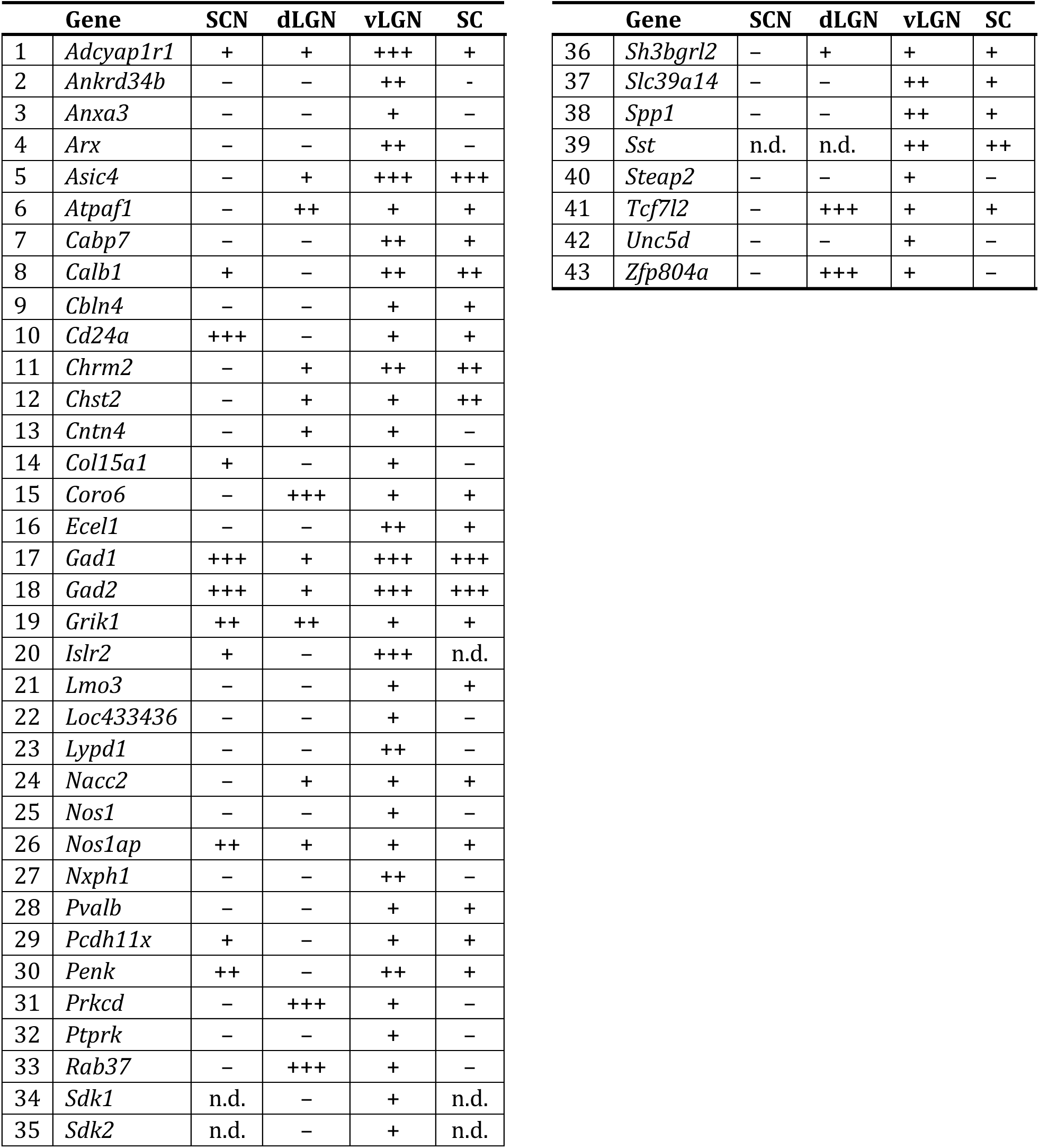
Riboprobe screen of genes enriched in vLGN. Symbols are qualitative indicators of expression in each region. The minus symbol indicates no cells expressing this mRNA observed and plus symbols indicate that cells were observed, ranging from some (+) to many (+++). SCN – suprachiasmatic nucleus; dLGN – dorsal lateral geniculate nucleus; vLGN – ventral lateral geniculate nucleus; SC – superior colliculus; n.d. – not done.

From this unbiased riboprobe screen, we identified two genes, *Nxph1* and *Arx*, whose expression in vLGN was restricted to cells in largely nonoverlapping domains. *Nxph1*, which encodes the α-Nrxn ligand Neurexophilin-1, was expressed in the most lateral portion of vLGN (Figure 1j-k). *Arx*, which encodes the homeobox transcription factor Aristaless Related Homeobox protein, was expressed in the most medial portion of vLGN (Figure 1l-m). Double-ISH, with *Gad1* riboprobes, revealed that both *Nxph1* and *Arx* mRNAs were generated by GABAergic cells in vLGN (Figure 1k,m,n-o). To test whether *Nxph1* and *Arx* marked GABAergic cell types in vLGNe and vLGNi respectively, we labeled retinal ganglion cell arbors in vLGN by intraocular injection of fluorescently conjugated Cholera Toxin Subunit B (CTB) (Figure 1p). Indeed, we found that *Nxph1*^+^ neurons reside in vLGNe and *Arx*^+^ neurons reside in vLGNi (Figure 1q). These results further suggested that transcriptionally distinct subsets of GABAergic neurons exist in vLGN and demonstrated cellular diversity in both laminae of vLGN – vLGNe and vLGNi (Figure 1r).

We next sought to determine whether GABAergic neurons residing within vLGNe or vLGNi could be further subdivided into distinct neuronal subtypes. Several of the genes identified as differentially enriched in the vLGN compared to dLGN were canonical markers of inhibitory interneurons in other parts of the brain (i.e. Somatostatin, Calbindin, Parvalbumin)(Figure 2a-c)(Tremblay *et al.* 2016; Lim *et al.* 2018). To label Somatostatin-expressing cells, we used two approaches: a *Sst-Cre::Rosa-Stop-tdT* reporter mouse in which all *Sst*^+^ cells were transgenically labeled with tdT and riboprobes against *Sst* mRNA. Both approaches revealed sparse *Sst*^+^ cells distributed broadly across both the vLGN and the intergeniculate leaflet (IGL), a slice of retinorecipient tissue separating vLGN from dLGN in mice (Figure 2d and data not shown). By coupling *Gad1* ISH in *Sst-Cre::Rosa-Stop-tdT* tissue, we found that >85% of *Sst*^+^ cells generated *Gad1* mRNA and were therefore GABAergic neurons. By labeling retinal axons with CTB and quantifying the density of tdT^+^ cells in vLGNe and vLGNi, we determined that *Sst*^+^ neurons were evenly distributed in vLGNe or vLGNi (Figure 2e-h). Next, we immunolabeled Calbindin-expressing neurons and found that Calb^+^ cells were in both IGL and vLGN, and most were also GABAergic (Figure 2i-k). When we labeled vLGNe and vLGNi with intravitreally injected CTB, we found that Calb^+^ cells were preferentially distributed in vLGNi, although they represent only a fraction of the cells in this region (Figure 2l-m). Finally, we labeled Parvalbumin-expressing neurons by generating a *Pvalb-Cre::Thy1-Stop-YFP* mouse in which all *Pvalb*^+^ cells were transgenically labeled with YFP (Figure 2n). We observed a modest number of *Pvalb*^+^ cells in vLGN, but unlike the broad distribution of *Sst^+^* and Calb^+^ neurons, *Pvalb^+^* cells were concentrated in what appeared to be a layer of vLGN and were absent from IGL (Figure 2n). By coupling *Gad1* ISH in *Pvalb-Cre::Thy1-Stop-YFP*, we found that >90% of *Pvalb*^+^ cells generated *Gad1* mRNA and were therefore GABAergic (Figure 2o-p). When we labeled retinal projections in vLGNe with CTB and *Pvalb^+^* GABAergic neurons by immunolabeling, we determined that *Pvalb^+^* cells were exclusively present in vLGNe (Figure 2q-r). It was noteworthy that *Sst^+^*, *Calb1^+^*, and *Pvalb^+^* cells were largely absent from the neighboring dLGN, suggesting that these GABAergic cell types were unique from previously studied dLGN interneurons.

**Figure 2.**
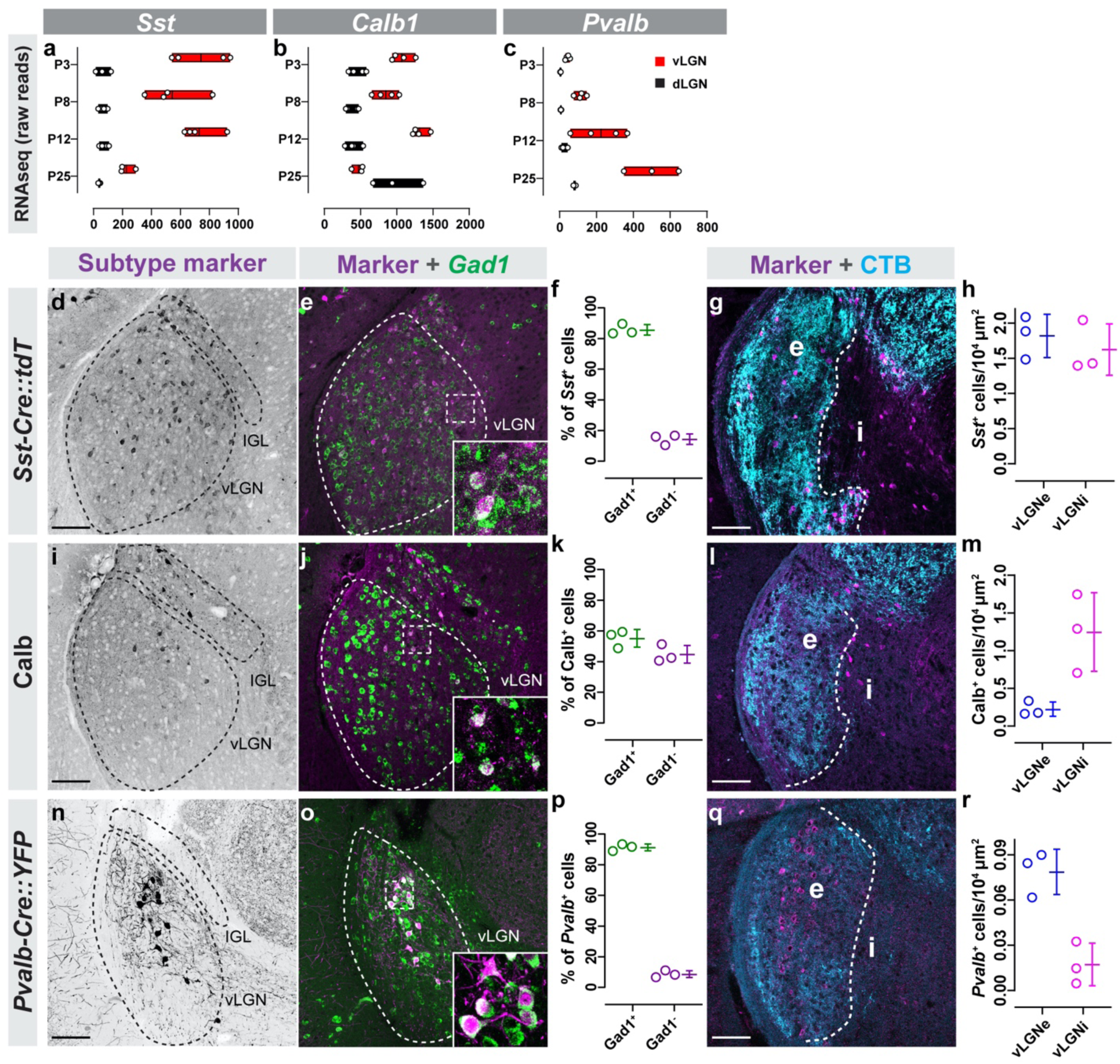
Canonical markers of GABAergic neurons label subsets of neurons in vLGN. (a-c) Raw transcript reads *Sst* (a)*, Calb1* (b)*, and Pvalb* (c) mRNA in vLGN and dLGN obtained by RNAseq. Individual datapoints plotted as white circles, min/max values are confined to the red bars, and vertical black line with bars depicts mean. (d) Transgenic labeling of *Sst^+^* neurons in vLGN of P60 *Sst-Cre::Rosa-Stop-tdT* mice. (e) ISH for *Gad1* mRNA in *Sst-Cre::Rosa-Stop-tdT* vLGN. (f) Quantification of the percentage of *Sst^+^* cells that co-express *Gad1* mRNA. Data points represent biological replicates, bars represent mean ± SD. (g) Transgenic labeling of *Sst^+^* neurons in vLGNe and vLGNi of *Sst-Cre:Rosa-Stop-tdT* mice following intravitreal CTB injection. (h) Quantification of the density of transgenically labeled *Sst^+^* cells in vLGNe and vLGNi. Data plotted as in (f). (i) Immunolabeling of Calb^+^ cells in P60 vLGN. (j) IHC-ISH for Calb protein and *Gad1* mRNA. (k) Quantification of Calb*+* and *Gad1*^*+*^ signal colocalization as seen in (j). Data plotted as in (f). (l) Calb*+* neurons in vLGNe and vLGNi visualized by Calb-immunolabeling and intravitreal CTB injection. (m) Quantification of the density of Calb-immunoreactive cells in vLGNe or vLGNi. Data plotted as in (f). (n) Transgenic labeling of *Pvalb^+^* neurons in P60 vLGN of P60 *Pvalb-Cre::Thy1-Stop-YFP* mice. (o) ISH for *Gad1* mRNA in *Pvalb-Cre::Thy1-Stop-YFP* vLGN. (p) Quantification of the percentage of *Pvalb^+^* cells that co-express *Gad1* mRNA. Data plotted as in (f). (q) Immunolabeling of Pvalb^+^ neurons in vLGNe and vLGNi of wildtype mouse following intravitreal CTB injection. (r) Quantification of the density of Pvalb^+^ cells density in vLGNe and vLGNi. Data plotted as in (f). All scale bars= 100 µm.

Since neither the *Pvalb*^+^ neurons (which preferred the vLGNe) nor Calb^+^ neurons (which preferred the vLGNi) labeled all of the cells in their respective subdomains, we hypothesized that an even richer heterogeneity of GABAergic cells existed. This led us to ask whether there were other types of GABAergic neurons which exhibited similar vLGN subdomain preferences. Using our riboprobe screen, we identified four additional genes that were generated by regionally restricted subsets of cells in vLGN: *Spp1*, *Penk*, *Lypd1*, and *Ecel1*. The transcripts for all of these genes were enriched in vLGN compared to dLGN (Figure 3a-d) and were generated by *Gad1*^*+*^ GABAergic neurons (Figure 3e-x).

**Figure 3.**
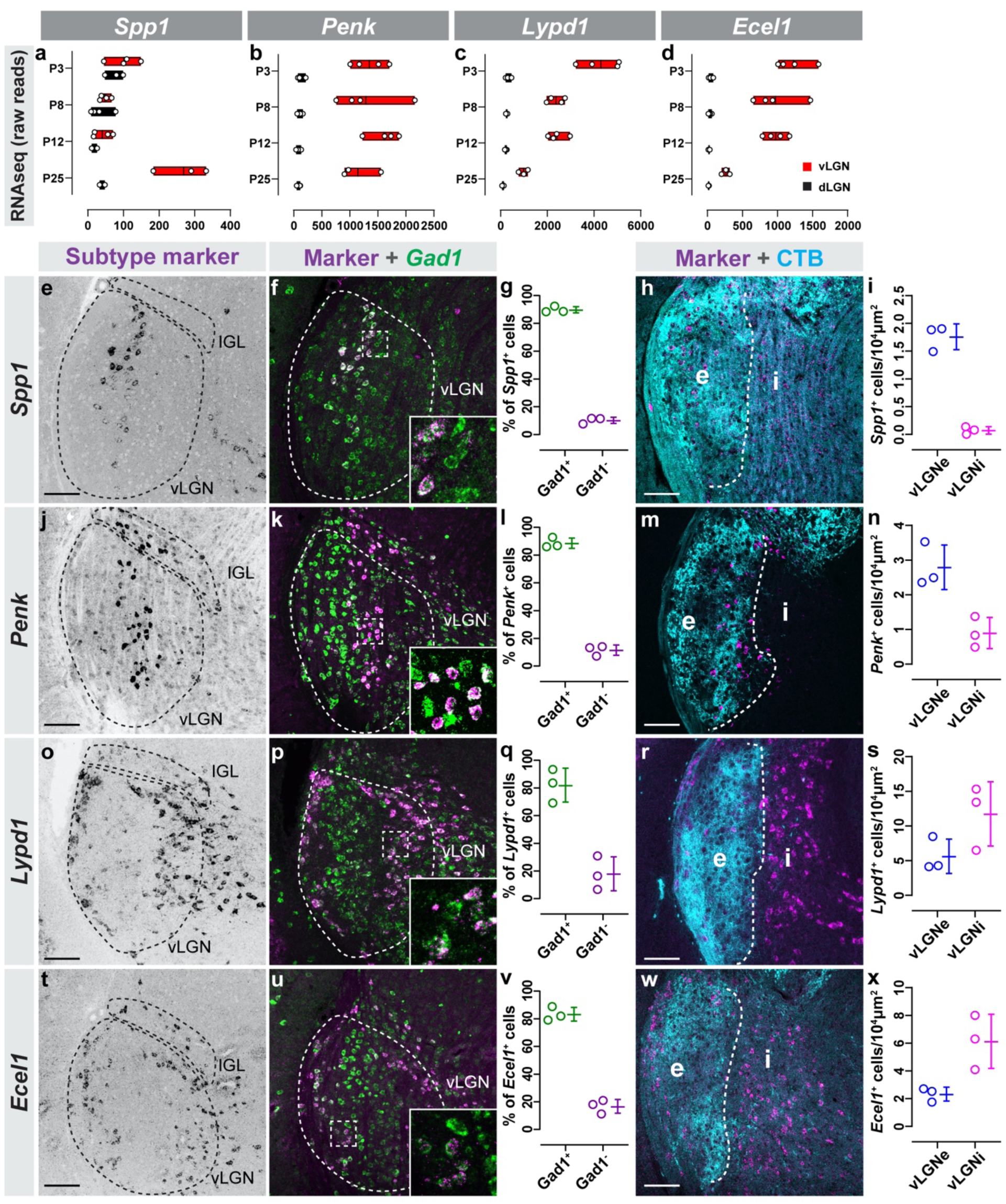
Novel genetic markers label subtypes of GABAergic neurons. (a-d) Raw transcript reads *Spp1* (a)*, Penk* (b)*, Lypd1* (c), and *Ecel1* (d) mRNA in vLGN and dLGN obtained by RNAseq. Individual datapoints plotted as white circles, min/max values are confined to the red bars, and vertical black line with bars depicts mean. (e) ISH for *Spp1* mRNA in P60 vLGN. (f) Double ISH for *Spp1* and *Gad1* mRNA in vLGN. (g) Quantification of the percentage of *Spp1+* cells that co-express *Gad1* mRNA. Data points represent biological replicates, bars represent mean ± SD. (h) ISH-labeling *Spp1+* neurons in vLGNe and vLGNi following intravitreal CTB injection. (i) Quantification of the density of *Spp1+* neurons in vLGNe and vLGNi. Data plotted as in (g). (j) ISH for *Penk* mRNA in P60 vLGN. (k) Double ISH for *Penk* and *Gad1* mRNA in vLGN. (l) Quantification of the percentage of *Penk^+^* cells that co-express *Gad1* mRNA. Data plotted as in (g). (m) ISH-labeling *Penk^+^* neurons in vLGNe and vLGNi following intravitreal CTB injection. (n) Quantification of the density of *Penk^+^* neurons in vLGNe and vLGNi. Data plotted as in (g). (o) ISH for *Lypd1* mRNA in P10 vLGN. (p) Double ISH for *Lypd1* and *Gad1* mRNA in vLGN. (q) Quantification of the percentage of *Lypd1+* cells that co-express *Gad1* mRNA. Data plotted as in (g). (r) ISH-labeling *Lypd1+* neurons in vLGNe and vLGNi following intravitreal CTB injection. (s) Quantification of the density of *Lypd1+* neurons in vLGNe and vLGNi. Data plotted as in (g). (t) ISH for *Ecel1* mRNA in P60 vLGN. (u) Double ISH for *Ecel1* and *Gad1* mRNA in vLGN. (v) Quantification of the percentage of *Ecel1+* cells that co-express *Gad1* mRNA. Data plotted as in (g). (w) ISH-labeling *Ecel1+* neurons in vLGNe and vLGNi following intravitreal CTB injection. (x) Quantification of the density of *Ecel1+* neurons in vLGNe and vLGNi. Data plotted as in (g). All scale bars = 100 µm.

Riboprobes generated against *Spp1*, which encodes the extracellular glycoprotein Osteopontin, revealed *Spp1*^+^ cells were sparsely present in the vLGN (and absent from both IGL and dLGN). Interestingly, *Spp1+* cells were distributed in a stratified fashion within vLGNe, just as we observed for *Pvalb^+^* cells (Figure 3e,h-i). ISH for *Penk*, which encodes Proenkephalin, revealed that *Penk* mRNA was present in a subset of vLGN cells and in many IGL neurons (Figure 3j-k). Like what we observed for *Spp1+* and *Pvalb^+^* neurons, *Penk^+^* neurons also appeared distributed in a stratified fashion. Labeling retinal afferents with CTB revealed that both *Spp1+* and *Penk*^+^ neurons were located in the retinorecipient vLGNe, although it was unclear if they were present in the same region (Figure 3m-n). Finally, ISH for *Lypd1*, which encodes LY6/PLAUR Domain Containing 1, and *Ecel1*, which encodes Endothelin Converting Enzyme Like 1, revealed that these genes exhibited similar cellular expression patterns in vLGN and IGL (Figure 3o,t). *Lypd1*^+^ and *Ecel1+* cells were sparsely distributed in the IGL and densely populated two distinct and separate regions of the vLGN, occupying both the lateral-most region of vLGNe and the entire vLGNi (Figure 3r-s,w-x). Based on similarities of expression patterns in both vLGN and IGL, it seemed likely that *Ecel1* and *Lypd1* mRNAs were generated in the same subsets of GABAergic neurons.

Taken together, these experiments reveal novel markers of transcriptomically and regionally distinct subsets of GABAergic neurons in vLGN. Importantly, not only did these cells have regional preferences, but *Pvalb*^+^, *Penk*^+^, *Spp1*^+^, *Ecel1*^+^, and *Lypd1*^+^ neurons each appeared to be organized into segregated strata that span the dorso-ventral axis of the vLGN.

### Transcriptomically distinct GABAergic neurons organize into adjacent sublaminae of vLGNe

We next asked whether these were in fact mutually exclusive groups of cells and whether they corresponded to distinct vLGNe sublaminae. We started by assessing whether *Spp1* and *Pvalb* mRNAs were generated by the same neurons or occupied the same dorso-ventral zone. Performing ISH on *Pvalb-Cre::Thy1-Stop-YFP* tissue revealed that *Spp1* mRNA was generated by some, but not all, *Pvalb^+^* cells (~50% *Spp1+* neurons were *Pvalb-*) and vice versa (~25% *Pvalb^+^* neurons were *Spp1*)(Figure 4a-d). *Spp1+Pvalb^+^*, *Spp^+^Pvalb-*, and *Pvalb-Spp1+* cells all appeared to reside within the same sublamina of vLGNe. To quantitatively assess cellular distribution in vLGN, we developed an automated script in ImageJ to measure fluorescent intensity along the medial-lateral axis of vLGN (Figure 4c’’). Fluorescent signals at each medial-lateral coordinate were averaged along the entire dorsoventral extent of vLGN (and between biological replicates) and the quantified data identified the same spatial region of vLGN as populated both *Spp1+* and *Pvalb^+^* cells (Figure 4e).

**Figure 4.**
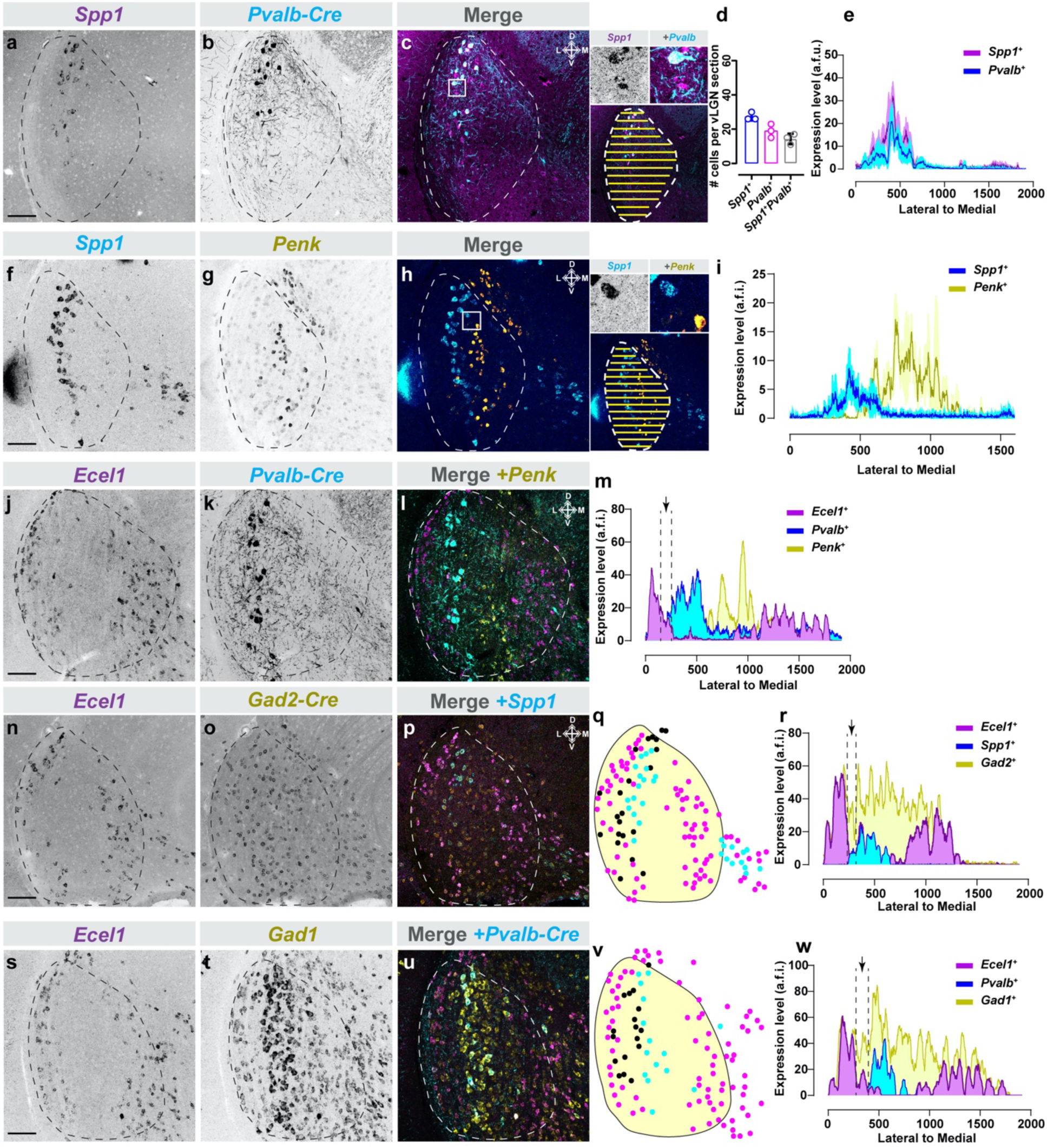
Transcriptomically distinct GABAergic neurons organize into discrete sublaminae of vLGNe. (a-c) ISH-labeling for Spp1+ neurons in vLGN of P60 *Pvalb-Cre::Thy1-Stop-YFP* transgenic reporter mice. Inset in (c) shows single channel and merged hi-magnification images and illustrates automated line scan approach used to quantify spatial expression. (d) Quantification of vLGN neurons which generate either or both *Spp1* and *Pvalb* mRNA. Data points represent biological replicates, bars represent mean ± SD. (e) Line scan analysis of spatial distribution of *Spp1+* and *Pvalb^+^* cells. Arbitrary fluorescence units (a.f.i.) are plotted against distance from the lateral-most part of the vLGN to the most medial. Solid line represents mean and shaded area represents SEM (n=3). (f-h) Double ISH for *Spp1* and *Penk* mRNA in P60 vLGN. Inset in (h) shows single channel and merged hi-magnification images and illustrates automated line scan approach used to quantify spatial expression. (i) Line scan analysis of spatial distribution of *Spp1+* and *Penk^+^* cells in vLGN plotted as in (e). (j-l) Double ISH for *Ecel1* and *Penk* mRNA in vLGN of P60 *Pvalb-Cre::Thy1-Stop-YFP* transgenic reporter mice. (m) Line scan analysis of spatial distribution of *Ecel1+*, *Penk^+^*, and *Pvalb^+^* cells in vLGN, plotted as in (e), revealing four discrete spatial domains of vLGN. The black arrow is pointing at a potential fifth layer between *Ecel1+* and *Pvalb^+^* layers. (n-p) Double ISH for *Ecel1* and *Spp1* mRNA in vLGN of P60 *Gad2-Cre::Sun1-Stop-GFP* transgenic reporter mice. (q) Schematic of subtype expression observed in (p), where each dot corresponds to a cell (magenta=*Ecel1+*, cyan=*Spp1+*, black=*Ecel1-Spp1-Gad2+*). Only *Gad2+* neurons between the two labeled layers are identified by black dots. (r) Line scan analysis of spatial distribution of *Ecel1+*, *Spp1+*, and *Gad2+* cells in vLGN, plotted as described in (e). Black arrow points at same region as arrow in (m). (s-u) Double ISH for *Ecel1* and *Gad1* mRNA in vLGN of P60 *Pvalb-Cre::Thy1-Stop-YFP* transgenic reporter mice. (v) Schematic of subtype expression observed in (u), where each dot corresponds to a cell (magenta=*Ecel1+*, cyan=*Pvalb^+^*, black=*Ecel1-Pvalb-Gad1^+^ neurons*), as in (q). (w) Line scan analysis of spatial distribution of *Ecel1+*, *Pvalb^+^*, and *Gad1*^*+*^ cells in vLGN, plotted as in (e). Black arrow points at same region as arrow in (m,r). All scale bars = 100 µm.

Next, we asked whether *Penk* was generated by either *Spp1+* or *Pvalb^+^* cells. For this, we used double ISH or genetic reporter lines. In both cases, we were unable to identify a single occurrence in which *Penk^+^* neurons co-expressed either *Pvalb* or *Spp1* (Figure 4f-h and data not shown). Moreover, these experiments clearly demonstrated that *Penk^+^* cells were not only transcriptomically distinct from *Spp1+* and *Pvalb^+^* cells, but also were present in an adjacent sublamina. Line scan analysis of fluorescence from *Penk* and *Spp1* double ISH experiments confirmed that these populations of GABAergic neurons were present in distinct sublaminae (Figure 4i). Next, we took a triple labeling approach (double ISH for *Ecel1*^+^ and *Penk*^+^ neurons in the transgenic *Pvalb-Cre::Thy1-Stop-YFP*) to test whether the sublaminae populated by *Pvalb^+^* and *Penk^+^* cells were distinct from those populated by *Ecel1+* cells (Figure 4j-m). Again, we observed no *Ecel1*^+^*Pvalb*^+^ or *Ecel1*^+^*Penk*^+^ neurons using this method, and quantitative analysis of *Ecel1/Penk/Pvalb* expression patterns along the medio-lateral extent of vLGN revealed at least three distinct sublaminae in vLGNe (Figure 4i).

Labeling of *Ecel1+*, *Pvalb^+^* and *Penk^+^* cells at once did not label all GABAergic cells in vLGN. In fact, it appeared as if the space between the lateral-most layer of *Ecel1*^+^ cells and the *Spp1*^+^ layer may represent another layer of GABAergic cells which we failed to identify with our riboprobe screen (arrow in Figure 4m). To test this idea, we performed two triple labeling experiments: one in which we labeled *Ecel1* and *Spp1* mRNA in *Gad2-Cre::Sun1-Stop-GFP* tissue, and one in which we labeled *Ecel1* and *Gad1* mRNAs in *Pvalb-Cre::Thy1-Stop-YFP* tissue (Figure 4n-w). In both cases, we identified GABAergic cells between the *Ecel1+* layer and the *Spp1+Pvalb^+^* layer. Line scan analysis confirmed a small population of *Gad2*^+^*Ecel1*-*Spp1*-neurons between the sublaminae containing *Ecel1*^+^ or *Spp1*^+^ neurons (Figure 4q-r,v-w), suggesting the existence of at least a fourth sublamina in vLGNe with yet-to-be-defined GABAergic neurons.

We recognized that our spatial registration of GABAergic subtypes into distinct sublaminae in vLGNe might have been an artifact of the coronal plane of section. To determine whether these sublaminae were in fact true structural components of vLGN, we performed axial (horizontal) sections of *Pvalb-Cre::Thy1-Stop-YFP* (Figure 5a-b). By performing double ISH on this tissue, we found that 1) *Spp1+* and *Pvalb^+^* cells reside in the same layer (and a population of cells express both transcripts), and 2) *Ecel1+*, *Pvalb^+^*, and *Penk^+^* cells populate distinct sublaminae of vLGNe (Figure 5c-k). Taken together, these data demonstrate, for the first time, that the vLGNe contains heterogeneous populations of transcriptomically distinct GABAergic cell types that map onto at least four adjacent sublaminae (that are not appreciable with conventional histochemical staining approaches).

**Figure 5.**
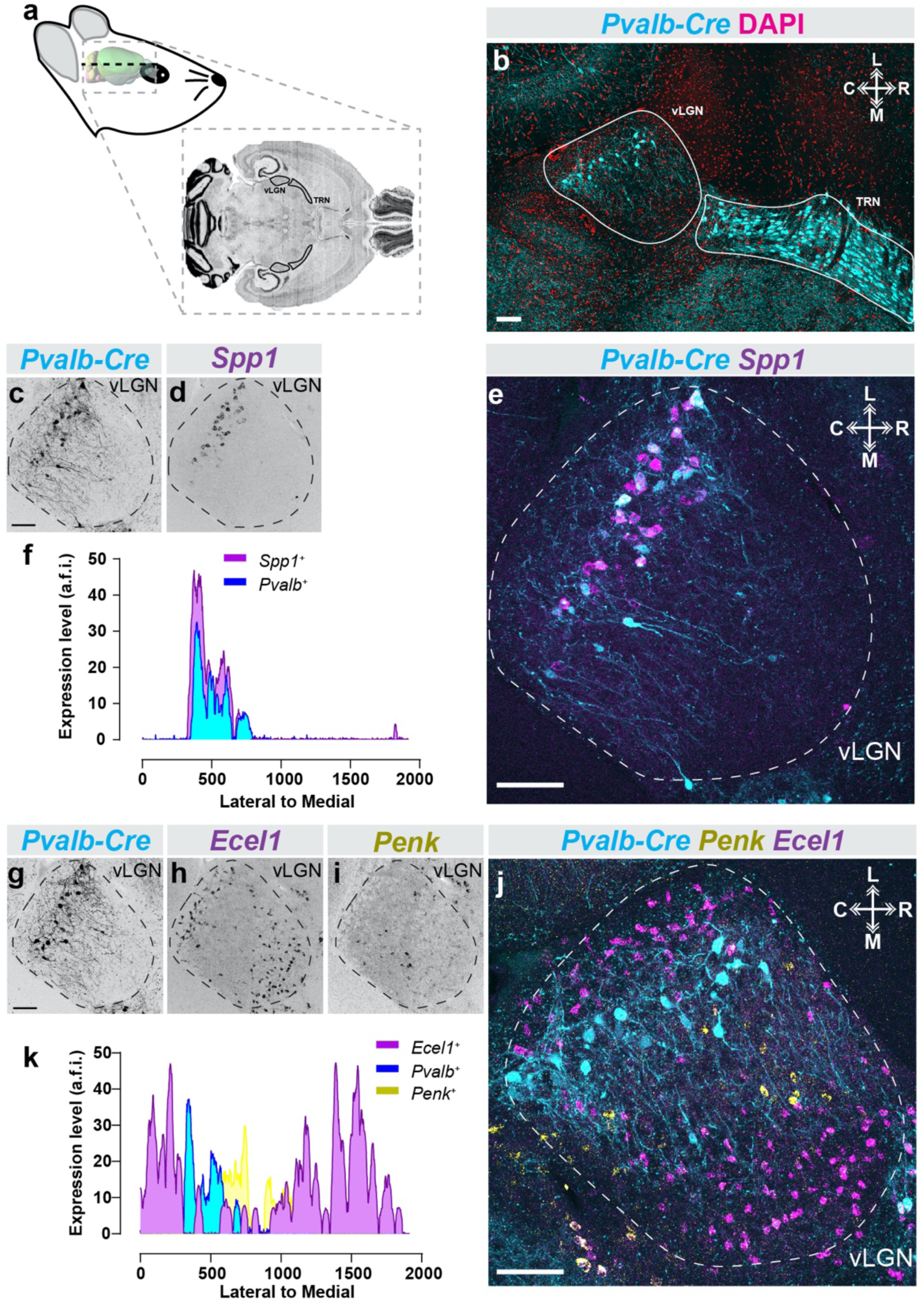
Subtype specific laminar organization of vLGN neurons along the rostro-caudal axis. (a) Schematic of horizontal mouse brain sectioning used to find vLGN. (b) Identification of the mouse vLGN and thalamic reticular nucleus (TRN) in the horizontal plane using the *Pvalb-Cre::Thy1-Stop-YFP* transgenic reporter and DAPI counter-staining. (c-e) ISH-labeling for *Spp1+* neurons in horizontal vLGN of P60 *Pvalb-Cre::Thy1-Stop-YFP* transgenic reporter mice. (f) Line scan analysis of spatial distribution of *Spp1+* and *Pvalb^+^* cells. Arbitrary fluorescence units (a.f.i.) are plotted against distance from the lateral-most part of the vLGN to the most medial. (g-j) Double ISH for *Ecel1* and *Penk* mRNA in horizontal vLGN of P60 *Pvalb-Cre::Thy1-Stop-YFP* transgenic reporter mice. (k) Line scan analysis of spatial distribution of *Ecel1+*, *Penk^+^*, and *Pvalb^+^* cells in vLGN, plotted as in (f), revealing four discrete spatial domains of vLGN. All scale bars = 100 µm.

### GABAergic neurons in the four sublaminae of vLGNe are retinorecipient

The laminar segregation of transcriptomically distinct cell types in vLGN raises the possibility that this organization functions to parse different streams of visual information. This led us to hypothesize that GABAergic cells in distinct sublaminae in vLGNe were directly innervated by retinal axons. In fact, while retinorecipient cells in dLGN are well defined, the retinorecipient neurons in vLGN are largely unknown. To anterogradely label retinorecipient cells in vLGN, we intravitreally injected a trans-synaptic adeno-associated virus expressing Cre recombinase (AAV2/1-hSyn-Cre-WPRE-hGH, referred to here as AAV1-Cre) into *Rosa-Stop-tdT* mice (Zingg *et al.* 2017)(Figure 6a-b). This trans-synaptic viral transfection strategy has previously been shown to accurately map monosynaptic connections and, in our hands, resulted in sparse tdT^+^ labeling of cells in retinorecipient brain regions (Figure 6B)(Zingg *et al.* 2017). Using this approach, we trans-synaptically labeled 53 retinorecipient neurons in vLGN (n=12 animals). Once we identified tdT^+^ cells in vLGN, we assessed their spatial localization relative to the sublaminae described above and performed ISH to determine whether the distinct subtypes of GABAergic neurons identified here were retinorecipient. We identified *Pvalb^+^*, *Spp1+*, *Ecel1+*, and *Penk^+^* cells in vLGNe that contained tdT, indicating that these populations of GABAergic neurons are capable of receiving monosynaptic input from the retina (Figure 6c-f).

**Figure 6.**
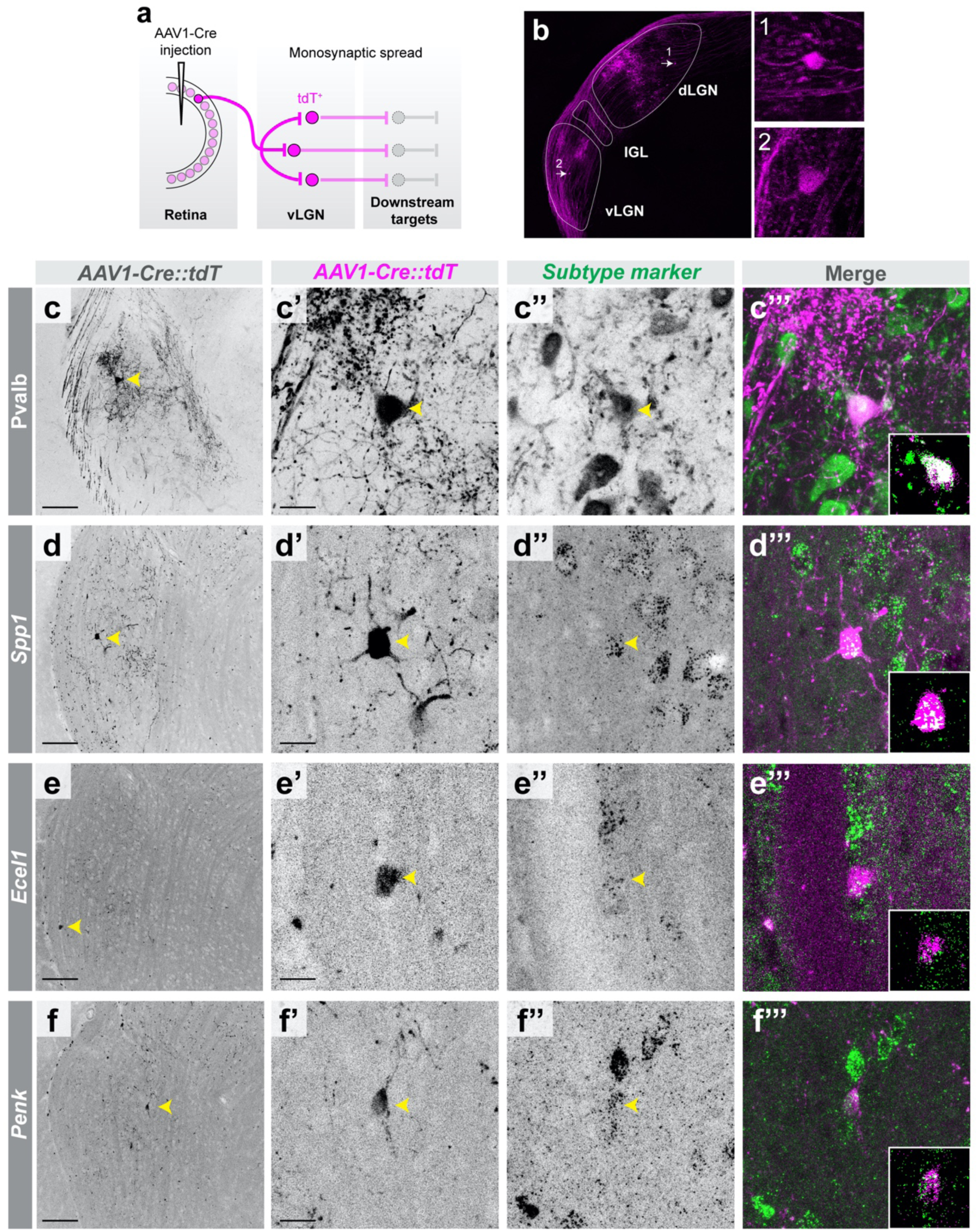
Several subtypes of GABAergic neurons in vLGNe receive direct retinal input. (a) Schematic of trans-synaptic viral tracing strategy to label retinorecipient vLGN neurons, where AAV1-Cre induces recombination (and therefore expression of tdT) in transfected cells. (b) Image of an LGN with retinorecipient cells labeled using strategy in (a). (c-c’) low-magnification and high-magnification micrographs, respectively, of labeled retinorecipient neuron in vLGN. (c’’-c’’’) Immunolabeling of Pvalb^+^ neurons in vLGN (c’’) and colocalization with tdT^+^ signal (c’’’). (d-d’) low-magnification and high-magnification micrographs, respectively, of labeled retinorecipient neuron in vLGN. (d’’-d’’’) ISH labeling of *Spp1+* neurons in vLGN (d’’) and colocalization with tdT^+^ signal (d’’’). (e-e’) low-magnification and high-magnification micrographs, respectively, of labeled retinorecipient neuron in vLGN. (e’’-e’’’) ISH labeling of *Ecel1+* neurons in vLGN (e’’) and colocalization with tdT^+^ signal (e’’’). (f-f’) low-magnification and high-magnification micrographs, respectively, of labeled retinorecipient neuron in vLGN. (f’’-f’’’) ISH labeling of *Penk^+^* neurons in vLGN (f’’) and colocalization with tdT^+^ signal (f’’’). Insets are single plane confocal images. Scale bars (c,d,e,f) = 100 µm, (c’,d’,e’,f’) = 25 µm.

We next sought to functionally confirm that some of these transcriptomically distinct cell types receive direct retinal input and to characterize their electrophysiological response properties. We utilized available transgenic reporter lines to record from and characterize several GABAergic subtypes in vLGN. Figure 7a-c provide examples of biocytin filled vLGN neurons recorded from *Pvalb-Cre::Thy1-Stop-YFP* (n=12), *Sst-Cre::Rosa-Stop-tdT* (n=15) and *GAD67-GFP* (n=5) mice along with representative voltage responses to current injection and synaptic responses evoked by optic tract (OT) stimulation. Overall, their dendritic architecture varied widely from each other. *Pvalb^+^* neurons had a hemispheric architecture with several elongated but sparsely branched primary dendrites, whereas *Sst^+^* neurons had much smaller bipolar dendritic fields that contained short multi-branched processes. Biocytin filled vLGN neurons from *GAD67-GFP* mice had similar morphologies to intrinsic interneurons in dLGN, displaying expansive and complex arbors that arise from opposite poles of fusiform-shaped soma (Guillery 1966; Charalambakis *et al.* 2019; Seabrook *et al.* 2013). The voltage responses to current injection also revealed differences in their intrinsic membrane and firing properties. For example, the resting membrane potential (Figure 7j) of *Sst^+^* (−57.5± 1.6 mV; *n* = 22) and *Pvalb^+^* (−59± 1.2 mV; *n* = 18) neurons was significantly (*P* <0.05, One-Way ANOVA) hyperpolarized compared to *GAD67-GFP^+^* neurons (−50 ± 2.2 mV; *n* = 6). The input resistance (Figure 7k) of *Pvalb^+^* (140± 18 MΩ; *n* = 18) was also significantly (*P* < 0.05, One-Way ANOVA) lower compared to *Sst^+^* (575± 46 MΩ; *n* = 22) and *GAD67-GFP^+^* neurons (278 ± 56 MΩ; *n* = 6). Responses to hyperpolarizing current pulses failed to evoke a large triangular shaped, rebound low threshold Ca2+ spike (**Figure7b,e,h**). However, both *Sst^+^* and *GAD67-GFP*^+^ neurons showed a strong depolarizing sag which is likely mediated by hyperpolarization activated cation conductance (Ih), while *Pvalb^+^* neurons showed little or no such inward rectification. For all three cell types, small depolarizing current steps (~80 pA) evoked a train of spikes that displayed spike frequency adaptation. However, when larger steps were employed (>100 pA), *Pvalb^+^* neurons displayed high frequency firing (84 ± 18 Hz). However, *Sst^+^* and *GAD67-GFP^+^* neurons had difficulty maintaining firing throughout the duration of the current step, showing a progressive inactivation of Na2+ spikes.

**Figure 7.**
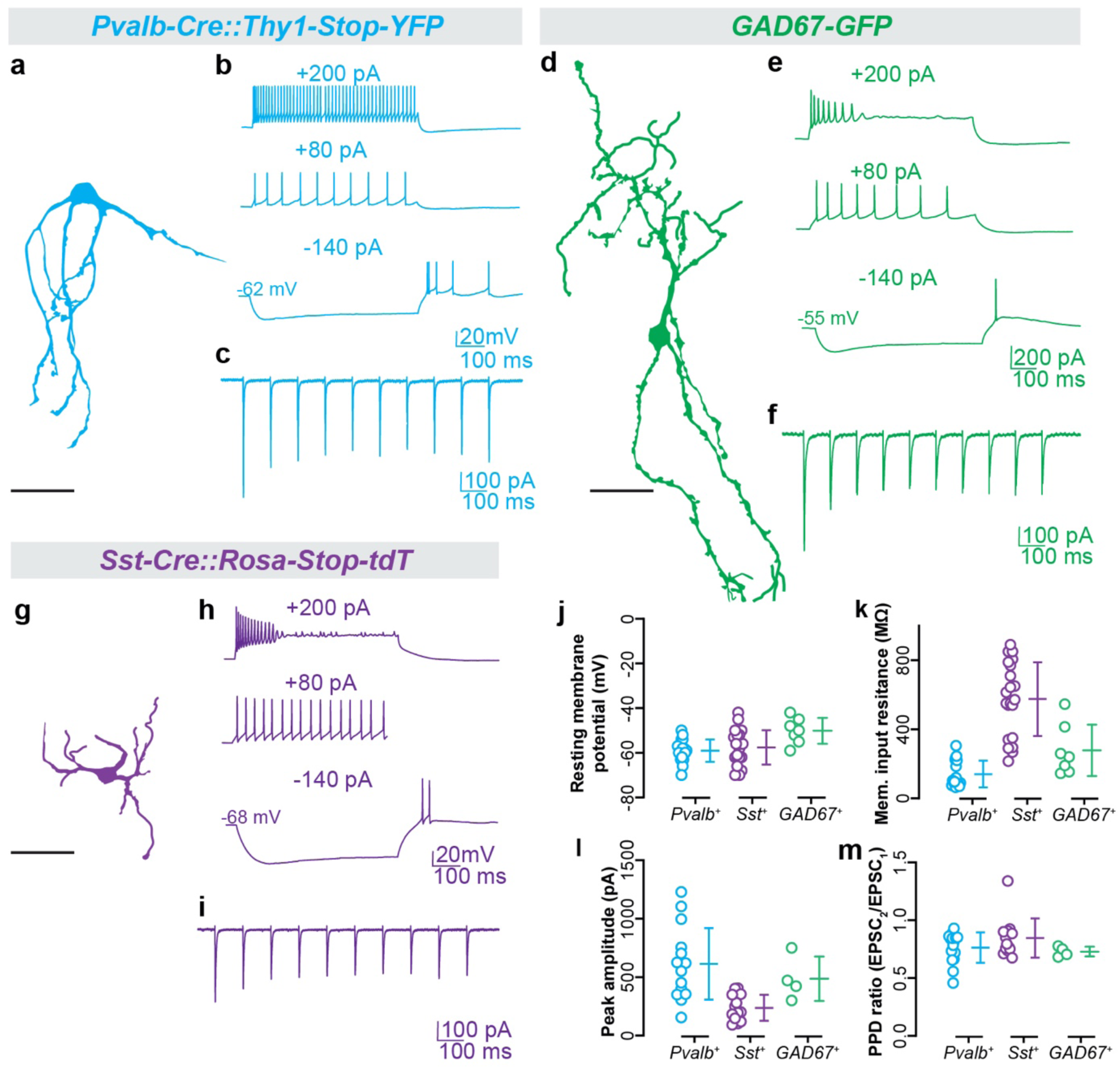
Morphology, synaptic responses, and membrane properties of GABAergic subtypes in vLGNe. (a-i) Representative confocal reconstructions of *Pvalb^+^* (a), *GAD67+* (d), and *Sst^+^* (g) neurons along with examples of their voltage responses to hyperpolarizing and depolarizing current pulses (top), and synaptic responses to optic tract (OT) stimulation (bottom). (j) Plot depicting resting membrane potential of *Pvalb^+^*, *Sst^+^*, and *GAD67+* neurons. (k) Plot depicting membrane input resistance of *Pvalb^+^*, *Sst^+^*, and *GAD67+* neurons. (l) Plot depicting peak excitatory postsynaptic currents (EPSC) amplitude of *Pvalb^+^*, *Sst^+^*, and *GAD67+* neurons. (m) Plot depicting paired pulse depression ration (PPD) during repeated stimulation of *Pvalb^+^*, *Sst^+^*, and *GAD67+* neurons. Each data point represents an individual value and bars reflect mean ± SEM. All scale bars = 50 µm. Gene SCN dLGN vLGN SC Gene SCN dLGN vLGN SC

Finally, we examined the synaptic responses that were evoked by OT stimulation (Figure 7c,f,i). As expected, OT stimulation evoked EPSC activity in neurons (38/46, 82.6%) located within vLGNe, while those recorded (n=14) in vLGNi failed to respond. The high prevalence of synaptic responses in the vLGNe was observed among *Pvalb^+^* (18/19), *Sst^+^* (13/15) and *GAD67-GFP^+^* (4/7) neurons. The peak amplitude of excitatory currents (Figure 7l) for *Pvalb^+^* (613± 79 pA; *n* = 15) and *GAD67-GFP^+^* (486± 95 pA; *n* = 4) neurons was significantly (*P* < 0.05, One-Way ANOVA) higher compared to *Sst^+^* neurons (237 ± 31; *n* =13). Synaptic responses evoked by a 10Hz stimulus train showed a moderate depression (75-85%) after the initial response. Measurements of paired pulse depression (Figure 7g) revealed no significant (p>0.1, One-way ANOVA) differences between *Pvalb^+^* (0.76± 0.03; *n* = 15), *Sst^+^* (0.84± 0.04; *n* = 13) and *GAD67-GFP^+^* (0.7± 0.02; *n* = 4) neurons.

## Discussion

In this study, we identified novel vLGN cell type markers which label GABAergic cells throughout the nucleus and distinguish it from its dorsal counterpart – the dLGN. By performing a series of multiplex labeling experiments using these newly identified markers, we revealed a remarkably organized laminar architecture of vLGNe, composed of at least four adjacent, transcriptionally distinct sublaminae. Using anterograde trans-synaptic viral tracing and patch-clamp electrophysiology, we determined that many of these regionally and transcriptomically distinct subtypes of GABAergic neurons receive direct retinal input. Thus, these data reveal a novel cellular organization of the vLGN and suggest such organization may have important implications for how different streams of retina-derived visual information are processed in this part of visual thalamus.

### Different types of hidden laminae exist in vLGN and dLGN

In contrast to the clear lamination of the primate and carnivore lateral geniculate nucleus, the rodent dLGN and vLGN have no obvious cytoarchitectonic lamination – save for the division of the vLGN into retinorecipient vLGNe and non-retinorecipient vLGNi (Niimi *et al.* 1963; Hickey & Spear 1976; Gabbott & Bacon 1994; Harrington 1997; Sabbagh *et al.* 2018). The neuronal cytoarchitecture of the rodent dLGN is composed of three well-defined classes of glutamatergic thalamocortical relay cells and just one or two classes of GABAergic inhibitory interneurons (roughly 10% of its neuronal population) (Arcelli *et al.* 1997; Jaubert-Miazza *et al.* 2005; Evangelio *et al.* 2018). While interneurons are present throughout the nucleus, relay cells of each class appear to exhibit regional preferences in their distribution within the dLGN (Krahe *et al.* 2011). These regional preferences in the rodent dLGN, however, do not alone capture the level of organization seen in primate and carnivore LGN. Instead, it appears that retinal afferents impart order in the rodent dLGN by segregating their arbors into “hidden laminae” both in terms of eye-specific domains and subtype-specific lamina (Martin 1986; Reese 1988; Hong & Chen 2011). Recent advances in transgenic labeling of individual RGC subtypes has revealed the precise architecture of these hidden laminae of subtype-specific retinal arbors in dLGN (Cruz-Martín *et al.* 2014; Huberman *et al.* 2008; Huberman *et al.* 2009; Kay *et al.* 2011; Kim *et al.* 2010; Kim *et al.* 2008; Martersteck *et al.* 2017; Rivlin-Etzion *et al.* 2011; Kerschensteiner & Guido 2017; Monavarfeshani *et al.* 2017). In this study, however, we identified a different kind of ‘hidden laminae’ within vLGN. Notably, the few identified subtypes of vLGN-projecting RGCs do not appear to segregate their terminal arbors into laminae in vLGN (with the notable exception that they are restricted to vLGNe) (Hattar *et al.* 2006; Osterhout *et al.* 2011; Monavarfeshani *et al.* 2017). The diffuse terminal arborization of these non-image forming subtypes of RGCs raises the question of whether visual information is, in fact, parsed into parallel channels in vLGN (Hattar *et al.* 2006; Osterhout *et al.* 2011). The stratification of transcriptomically distinct neurons presented in this study into adjacent sublaminae in vLGNe may contribute to the parallel processing of sensory information in this brain region. Just as the organization of retinal inputs imposes order on the otherwise less organized cytoarchitecture of dLGN, the diversity and organization of intrinsic cells in vLGN may impose order on the unorganized input it receives from the retina.

### Do laminated retinorecipient circuits organize visual pathways through the vLGN?

Is the quantifiable segregation of distinct GABAergic subtypes into sublaminae a potential means of organizing visual information arriving from the retina? To address this, we sought to determine whether these subtypes were directly innervated by RGCs. Using anterograde trans-synaptic tracing, we identified *Ecel1+*, *Pvalb^+^*, *Spp1+*, and *Penk^+^* cells as retinorecipient, together representing at least three sublaminae of vLGNe. While we failed to observe any retinorecipient cells in vLGNi using this method, it certainly remains possible that the dendrites of cells in vLGNi extend into vLGNe and receive monosynaptic input from the retina. Nevertheless, our viral tracing results suggest that the cell type-specific organization of the vLGN is relevant to organizing visual input. Such organization of retinorecipient cells hints at a potentially novel role for vLGN in visual processing, by which incoming visual input is sampled by specific GABAergic cell types and parsed into parallel channels of sensory information to be transmitted to downstream targets.

The specificity of these organized cell types in vLGN raises questions of whether they are projection neurons or local interneurons. Our electrophysiological and morphological analyses of some of these cell types suggests that cells labeled in *GAD67-GFP* mice are likely vLGN interneurons. Based on their preponderance in the vLGN, it is likely that at least some of the other subtypes of GABAergic neurons here are projection neurons. Unlike the dLGN, which has afferent projections to just visual cortex and the thalamic reticular nucleus, neurons in vLGN project to a diverse set of over a dozen downstream subcortical regions including the SC, the nucleus of the optic tract and other pretectal nuclei (Cadusseau & Roger 1991; Swanson *et al.* 1974; Trejo & Cicerone 1984), the suprachiasmatic nucleus (at least in hamsters)(Harrington 1997), the habenula (Oh *et al.* 2014; Huang *et al.* 2019), and zona incerta (Ribak & Peters 1975), all contributing to visuomotor, ocular, vestibular, circadian, and other innate behaviors (Monavarfeshani *et al.* 2017). It is known that some GABAergic cells in vLGN receive retinal input and project to the lateral habenula, although it remains unclear which GABAergic subtypes this includes (Huang *et al.* 2019).

*Might the different transcriptionally distinct GABAergic cell types in vLGN each project to different downstream nuclei and contribute to unique functions and behaviours?* First, we note here that it is conceivable that vLGN not only organizes and transmits visual information in separate channels, but also samples specific features of retinal input in sublamina-specific manner – consistent with labeled-line theory. Unfortunately, we do not yet have subtype specific resolution of vLGN projection neurons but hope that the data presented here will help to create a molecular toolkit for such circuit tracing in future studies. Such experiments will be crucial in a) determining whether parsing visual information into these hidden laminae is important for parallel processing and b) whether there is a functional and/or projection-specific logic to the lamination of vLGN cell types.

### Transcriptomic heterogeneity underlying cellular diversity in vLGN

There is a current push to use unbiased approaches to identify all of the cell types in the brain – essentially to create a ‘parts’ list. These studies have typically employed single cell RNA sequencing (scRNAseq) to understand the development, structure, and evolution of the brain (Krienen *et al.* 2019; Saunders *et al.* 2018; Peng *et al.* 2019). In a neuroscience community that consists of ‘lumpers’ and ‘splitters’, it is clear we are currently in an era of ‘splitting’ – as our technology to detect transcriptomic heterogeneity of cell types evolves, so too does the granularity of their discretization. Here, we have begun to ‘split’ the rodent vLGN into many distinct GABAergic cell types.

Rather than performing scRNAseq, we tackled the heterogeneity question by using bulk RNAseq and generating riboprobes with no *a priori* knowledge to perform a battery of *in situ* hybridizations for transcriptional heterogeneity in vLGN. By generating riboprobes against single molecules, we delineated distinct populations of transcriptomically and spatially distinct cells, an advantage of this approach over scRNAseq. Conservatively, our results demonstrate the existence of at least a half dozen discrete and separable subtypes of GABAergic neurons in vLGN, (although the number is likely much higher than this) and four distinct, adjacent sublaminae of GABAergic neurons in vLGNe.

Whether there are six GABAergic cell types in vLGN or many more has made us ponder the old question (Cajal 1893) of *how does one define a cell type?* Traditionally, classes and types of neurons have been characterized on the basis of morphological, electrophysiological, neurochemical, connectomic, or genetic information (Sanes & Masland 2015). Unfortunately, rarely are all these aspects of neuronal identity accounted for in a comprehensive way to glean a more accurate understanding about the structure of the nervous system. This has led to discrepant subtype classification across technical methodologies and challenges to comparing results between research groups. Here, we used spatial distribution and transcriptional profiles to classify neurons into distinct subtypes. Molecular markers remain at present a leading characteristic for such classifications, though they are not without their limitations. It may well be that vLGN neurons can be further subdivided if classified by 2-3 molecular markers rather than one (as we observed in *Spp1+* and *Pvalb^+^* neurons). However, the very fact that differential expression of one molecule was sufficient to differentiate two vLGN populations is a strong indicator that the nucleus as a whole is more diversely populated than previously appreciated. Nevertheless, we acknowledge here the efforts to create a more comprehensive framework to classifying cell types in the field. A set of recent studies represent a major step towards this goal by utilizing Patch-seq to simultaneously characterize cortical GABAergic neurons electrophysiologically, morphologically, and transcriptomically (Scala *et al.* 2020; Gouwens *et al.* 2020). Our approach here did not take into account these additional functional aspects of neuronal identity for all of the GABAergic cell types identified.

Our data, when taken together, suggest the possibility that functional organization of non-image-forming information from retina to vLGN is extracted from the segregation of transcriptionally distinct retinorecipient cells. We view these results as a framework for further dissecting the structure, circuitry, and functions of the vLGN at a cell-type specific level. How this heterogeneity and organization contributes to the yet-to-be determined functions of the vLGN remains to be defined.

## Acknowledgments

This work was supported in part by the following grants from the National Institutes of Health: EY021222 (M.A.F), EY030568 (M.A.F), NS105141 (M.A.F.), NS113459 (U.S.), and EY012716 (W.G.). The AAV1-Cre virus (pENN.AAV.hSyn.Cre.WPRE.hGH) was a gift from J.M. Wilson. We thank Dr. A.S. LaMantia (Fralin Biomedical Research Institute, Virginia Tech) for thoughtful comments on the manuscript.

## Abbreviations

RGC: retinal ganglion cells
LGN: lateral geniculate nucleus
dLGN: dorsal lateral geniculate nucleus
vLGN: ventral lateral geniculate nucleus
IGL: intergeniculate leaflet
SC: superior colliculus
SCN: suprachiasmatic nucleus
IHC: immunohistochemistry
ISH: in situ hybridization
AAV: adeno-associated virus
RRID: research resource identifier

